# Nanotrap^®^ particles improve detection of SARS-CoV-2 for pooled sample methods, extraction-free saliva methods, and extraction-free transport medium methods

**DOI:** 10.1101/2020.06.25.172510

**Authors:** RA Barclay, I Akhrymuk, A Patnaik, V Callahan, C Lehman, P Andersen, R Barbero, S Barksdale, R Dunlap, D Goldfarb, T Jones-Roe, R Kelly, B Kim, S Miao, A Munns, D Munns, S Patel, E Porter, R Ramsey, S Sahoo, O Swahn, J Warsh, K Kehn-Hall, B Lepene

## Abstract

Here we present a rapid and versatile method for capturing and concentrating SARS-CoV-2 from transport medium and saliva using affinity-capture magnetic hydrogel particles. We demonstrate that the method concentrates virus prior to RNA extraction, thus significantly improving detection of the virus using a real-time RT-PCR assay across a range of viral titers, from 100 to 1,000,000 viral copies/mL; in particular, detection of virus in low viral load samples is enhanced when using the method coupled with the IDT 2019-nCoV CDC EUA Kit. This method is compatible with commercially available nucleic acid extraction kits, as well with a simple heat and detergent method. Using transport medium diagnostic remnant samples that previously had been tested for SARS-CoV-2 using either the Abbott RealTime SARS-CoV-2 EUA Test (n=14) or the Cepheid Xpert Xpress SARS-CoV-2 EUA Test (n=35), we demonstrate that our method not only correctly identifies all positive samples (n = 17) but also significantly improves detection of the virus in low viral load samples. The average improvement in cycle threshold (Ct) value as measured with the IDT 2019-nCoV CDC EUA Kit was 3.1; n = 10. Finally, to demonstrate that the method could potentially be used to enable pooled testing, we spiked infectious virus or a confirmed positive diagnostic remnant sample into 5 mL and 10 mL of negative transport medium and observed significant improvement in the detection of the virus from those larger sample volumes.

## Introduction

Severe acute respiratory syndrome coronavirus 2 (SARS-CoV-2) is a member of the Coronaviridae family and is responsible for the pandemic outbreak of the coronavirus disease 2019 (COVID-19) that emerged in December 2019 in Wuhan city, Hubei Province, China ^1^. The disease is characterized by fever, dry cough, fatigue, anorexia, shortness of breath and myalgia ^2, 3^. COVID-19 rapidly spread and by March 11, 2020, the World Health Organization (WHO) declared COVID-19 as a global pandemic ^4^. As of June 7, 2020, there have been more than 7 million COVID-19 cases and 411,177 deaths worldwide ^5^. Such a fast-acting and massive outbreak throughout the world has led to severe impacts on the health care systems and economies of countries around the globe.

The rapid spread of the virus led to a tremendous increase in demand for COVID-19 diagnostic testing worldwide. At present, the United States Centers for Disease Control and Prevention (CDC) recommends diagnosis of acute SARS-CoV-2 infection via measurement of viral nucleic acid and antigen tests (https://www.cdc.gov/coronavirus/2019-nCoV/hcp/clinical-criteria.html). Real-time reverse transcriptase-polymerase chain reaction (real-time RT-PCR) is a reliable and relatively fast method for the identification of pathogenic nucleic acids in patient samples ^6^. However, it has several disadvantages. High quality RNA is critical to real-time RT-PCR assays. Thus, the initial step often involves purification of SARS-CoV-2 RNA from patient samples using commercial RNA purification kits, which have multiple steps and are time consuming. Moreover, in recent months, the manufacturers of these kits have struggled to keep up with demand, and there have been reports of shortages in the United States ^7^. Another disadvantage is the requirement for relatively high concentrations of the genetic material in a sample. Although real-time RT-PCR is considered to be a sensitive method for the detection of nucleic acid, the limit of detection for SARS-CoV-2 RNA is reported to be between 200 to 77,440 copies/mL, depending on the RNA extraction method and real-time RT-PCR assay being used ^6, 8, 9^.

We sought to address these issues with a method, based on affinity-capture hydrogel particles (called Nanotrap® particles), that captures and concentrates SARS-CoV-2 from samples to improve the detection of the virus when used with CDC-recommended SARS-CoV-2 assays. Here, we demonstrate that a 5-minute Nanotrap particle capture step significantly increases the sensitivity of SARS-CoV-2 real-time RT-PCR assays when used in conjunction with commercial RNA extraction kits. We also show that Nanotrap particles significantly improve sensitivity of SARS-CoV-2 real-time RT-PCR assays when used with simple heat and detergent extraction methods, in both saliva and transport medium samples. Furthermore, with this method, we identified viral RNA in several diagnostic remnant samples that previously had tested negative for SARS-CoV-2. Finally, we tested and confirmed the ability of a Nanotrap particle method to improve detection of SARS-CoV-2 RNA in pooled patient sample mimics, an approach which is a promising way forward for addressing the massive testing scale-ups that are necessary. Taken together, the methods here are quick and easy to implement, requiring only a magnetic tube rack to separate the particles and captured viruses, and they significantly improve the detection capability of current SARS-CoV-2 test strategies.

## Methods and Materials

### Cells, Viruses, and Reagents

Vero E6 cells were obtained from ATCC (CRL-1586) and were grown in DMEM complete medium, consisting of 10% FBS, 5% L-glutamine, and 5% penicillin/streptomycin, at 37°C and 5% CO_2_. Infectious, replication-competent and heat-inactivated SARS-CoV-2 were obtained from BEI Resources (NR-52281 and NR-52286, respectively). For transport medium experiments, Puritan UniTranz®-RT Universal Transport Medium (UT-300) was utilized. Nanotrap Magnetic Virus Particles (SKU 44202) were provided by Ceres Nanosciences Inc. Dulbecco’s phosphate-buffered Saline Solution without Ca2+ and Mg2+ (PBS) was used during viral extraction. TRIzol™LS was purchased from ThermoFisher Scientific (Cat. #10296010). Patient pooled saliva was purchased from BIOIVT (Saliva-1902492).

### Diagnostic Remnant Samples

Fourteen diagnostic remnant samples of patient swabs in viral transport media were purchased from Discovery Life Sciences. Each sample previously had been tested for SARS-CoV-2 using the Abbott RealTime SARS-CoV-2 test. Nine samples (sample ID’s 101-109) previously had tested positive. Five samples (sample ID’s 110-114) had previously tested negative. An additional thirty-five diagnostic remnant samples were obtained from Hancock Regional Hospital in Greenwood, IN, all of which had been previously tested for SARS-CoV-2 using the Cepheid Xpert® Xpress SARS-CoV-2 test. Eight samples (sample ID’s 201-208) had previously tested positive for COVID-19. Twenty-seven had previously tested negative (sample ID’s 209-235). Most of the samples were frozen after testing at the original sites and were shipped frozen. Due to challenges associated with shipping, a few of the samples were never frozen and were shipped and received refrigerated and at least one sample underwent multiple freeze-thaw cycles. All frozen samples were thawed directly prior to processing.

### Concentrating SARS-CoV-2 from Transport Medium

SARS-CoV-2 (heat-inactivated or infectious, replication-competent) was spiked into Puritan Universal Transport Medium at various concentrations. Two hundred microliters (1 mg) of Nanotrap particles in the storage solution were added to 1.5 mL microcentrifuge tubes and pulled out of solution using a DynaMag™-2 magnet from ThermoFisher (12321D). The supernatant was removed, and one milliliter of spiked transport medium was added to the particles; a quick mixing of the sample and the Nanotrap particles was performed using a micropipette. For large volume samples (5 or 10 mL of spiked transport medium), three hundred microliters (1.5 mg) of Nanotrap particles were used. Unless otherwise described, samples were incubated with Nanotrap particles at room temperature for 5 minutes to capture virus. Other than the initial mixing of the sample and the particles, no mixing was required during the 5-minute incubation. For incubations longer than 5 minutes, samples were inverted once every 5 minutes. Following incubation, Nanotrap particles were pulled out of solution with the DynaMag-2 magnet and the supernatant was discarded.

### Concentrating SARS-CoV-2 from Saliva

Heat-inactivated SARS-CoV-2 was spiked into the saliva at various concentrations. The saliva was allowed to sit for 3 minutes while aggregates settled. The liquid phase of saliva, which contained the virus, was moved into a new tube and was diluted in PBS with 0.05% Tween-20 (one-part saliva was added to two-parts PBS/Tween). Aliquots of 1.8 mL of diluted saliva were added to three hundred microliters (1.5 mg) of Nanotrap particles, which previously had been removed from their storage solution using a DynaMag-2 magnetic rack. Samples were incubated at room temperature for 10 minutes (at the 5-minute mark, samples were inverted once). Following incubation, Nanotrap particles were pulled out of solution with a DynaMag-2 magnet and the supernatant was discarded.

### Nucleic Acid Extraction

We evaluated the impact of using Nanotrap particles to capture and concentrate SARS-CoV-2 on several nucleic acid extraction methods: the QIAamp® Viral RNA Mini Kit from QIAGEN (52906); the RNeasy® Mini Kit from QIAGEN (74106); a TRIzol LS method used in combination with the RNeasy Mini Kit; a heat extraction method, which used a short heating step to lyse the virus; and a heat and detergent method, which used a short heating step and some detergent to lyse the virus. Those extraction methods are described below in their uses with and without Nanotrap particles.

When Nanotrap particles were used to capture and concentrate virus prior to nucleic acid extraction, nucleic acids were extracted under the following conditions. For the QIAamp Viral RNA Mini Kit, the Nanotrap particle pellet was resuspended in 150 µL of PBS and 560 µL of viral lysis buffer (Buffer AVL) was added to the re-suspended particles and allowed to incubate for 10 minutes at room temperature. Following this, particles were pulled out of solution using the DynaMag-2 magnet, and the supernatant was collected and added to 560 µL of ethanol, upon which the manufacturer’s protocol was followed without any deviations.

For the TRIzol LS method, the Nanotrap particle pellet was resuspended in 100 µL of water, mixed with 300 µL of TRIzol LS for virus inactivation, and incubated for 10 minutes at room temperature. This was followed by mixing with 200 µL of chloroform, spinning down, collecting the upper phase, mixing the upper phase with 300 µL of RLT buffer from the RNeasy Mini Kit, and adding an equal volume of 70% ethanol to the mixture. The next purification steps were followed according to the manufacturer’s protocol for the RNeasy Mini Kit.

For the direct extraction method, the Nanotrap particle pellet was resuspended in a detergent buffer and was incubated for 5 minutes at 95°C. For heat only extraction, the Nanotrap particle pellet was resuspended in 50 µL of water and heated for 10 minutes at 95°C. Following this, the particles were pulled out of solution using the DynaMag-2 magnet, and the supernatant was collected for analysis. At the end of each of these methods, the resulting solution, with extracted RNA, was loaded into the downstream assay.

In samples processed without Nanotrap particles, nucleic acids were extracted under the following conditions. For the QIAamp Viral RNA Mini Kit, nucleic acids were extracted from 150 µL of sample according to the manufacturers’ protocol.

For the RNeasy Mini Kit, nucleic acids were extracted from 150 µL of sample according to the manufacturers’ protocol.

For the TRIzol LS method, 100 µL of sample was mixed with 300 µL of TRIzol LS for virus inactivation. After that samples were mixed with 200 µL of chloroform, spun down, the upper phase was mixed with 300 µL RLT buffer from the RNeasy Mini Kit, and an equal volume of 70% ethanol was added. This was followed by loading samples onto the column and RNA was purified according to the RNeasy Mini Kit recommendations.

For the direct extraction method, 50 µL of sample was heated at 95°C for ten minutes. At the end of each of these extraction methods, the resulting solution, with extracted RNA, was loaded into the downstream assay.

### Real-Time RT-PCR

For heat-inactivated SARS-CoV-2 samples, two assays, the Primerdesign Ltd COVID-19 Genesig Real-Time CE-IVD/EUA PCR assay (Z-COVID-19) and the IDT 2019 nCoV CDC EUA Kit (1006770), were used for real-time RT-PCR. Per Primerdesign’s instructions, 10 µL of the Oasig™OneStep 2X qRT-PCR Master Mix, 2 µL of COVID-19 primer/probe, and 8 µL of RNA template were added to each well. PCR conditions were performed according to Primerdesign’s instructions on a Roche LightCycler^®^ 96. All samples were run in technical triplicate. Resulting Ct values were averaged across replicates and standard deviations were calculated.

Per IDT’s recommendation, TaqPath™1-Step RT-qPCR Master Mix from ThermoFisher (A15300) was used in the IDT 2019 nCoV CDC EUA assay. Per IDT’s instructions, each PCR reaction used 8.5 µL of nuclease free water, 5 µL of the TaqPath solution, 1.5 µL of the N1 primer/probe, and 5 µL of RNA template. PCR conditions were performed according to IDT’s instructions on a Roche LightCycler 96. All samples were run in technical triplicate. Resulting Ct values were averaged across replicates and standard deviations were calculated. All heat-inactivated SARS-CoV-2 experiments utilized this assay unless otherwise denoted. All patient remnant samples utilized this assay.

For infectious, replication-competent SARS-CoV-2 samples, we used the N1 primer from the IDT 2019 nCoV CDC EUA Kit (1006770). For the real-time RT-PCR assay, we used the RNA UltraSense™ One-Step Quantitative RT-PCR System from Applied Biosystems™ (11732927) according to manufacturer’s recommendation. Each PCR reaction contained 1 µL UltraSense Enzyme Mix, 5 µL RNA UltraSense 5X Reaction Mix, 1.5 µL of primer/probe mixture from IDT, 0.4 µL ROX Reference Dye, 8.1 µL of nuclease free water, and 5 µL of RNA template. The real-time RT-PCR was performed using an ABI StepOne Plus instrument. All samples were run in biological triplicate unless otherwise described.

## Results

### Nanotrap^®^ Particles Capture and Concentrate Inactivated SARS-CoV-2 from Transport Medium and Saliva

In previous studies, we have shown that hydrogel particles functionalized with high-affinity chemical baits are able to capture and concentrate from biological samples a variety of respiratory viruses, including Influenza A and B, RSV, and Coronavirus 229-E ^10-13^. In light of this, we asked whether the same particles could capture and concentrate SARS-CoV-2 from transport medium and saliva in order to improve diagnostic testing for COVID-19.

In all experiments, a simple four-step workflow was used to capture and concentrate the virus using hydrogel particles made by Ceres Nanosciences (Nanotrap particles). Briefly, viral transport medium spiked with heat-inactivated SARS-CoV-2 was mixed with Nanotrap particles and incubated at room temperature. The Nanotrap particles and the captured viruses were separated from the solution using a magnet and the supernatant was removed. The particle pellet was re-suspended in lysis buffer, which lysed the virus and released its nucleic acid. The Nanotrap particles were pelleted a second time with a magnet, and the nucleic acid-containing supernatant was ready for analysis by real-time RT-PCR (if using a direct nucleic acid extraction method, **Fig. 1A**) or for further RNA purification (if using a commercial nucleic acid kit extraction method, **Fig. 1B**).

**Figure 1:**
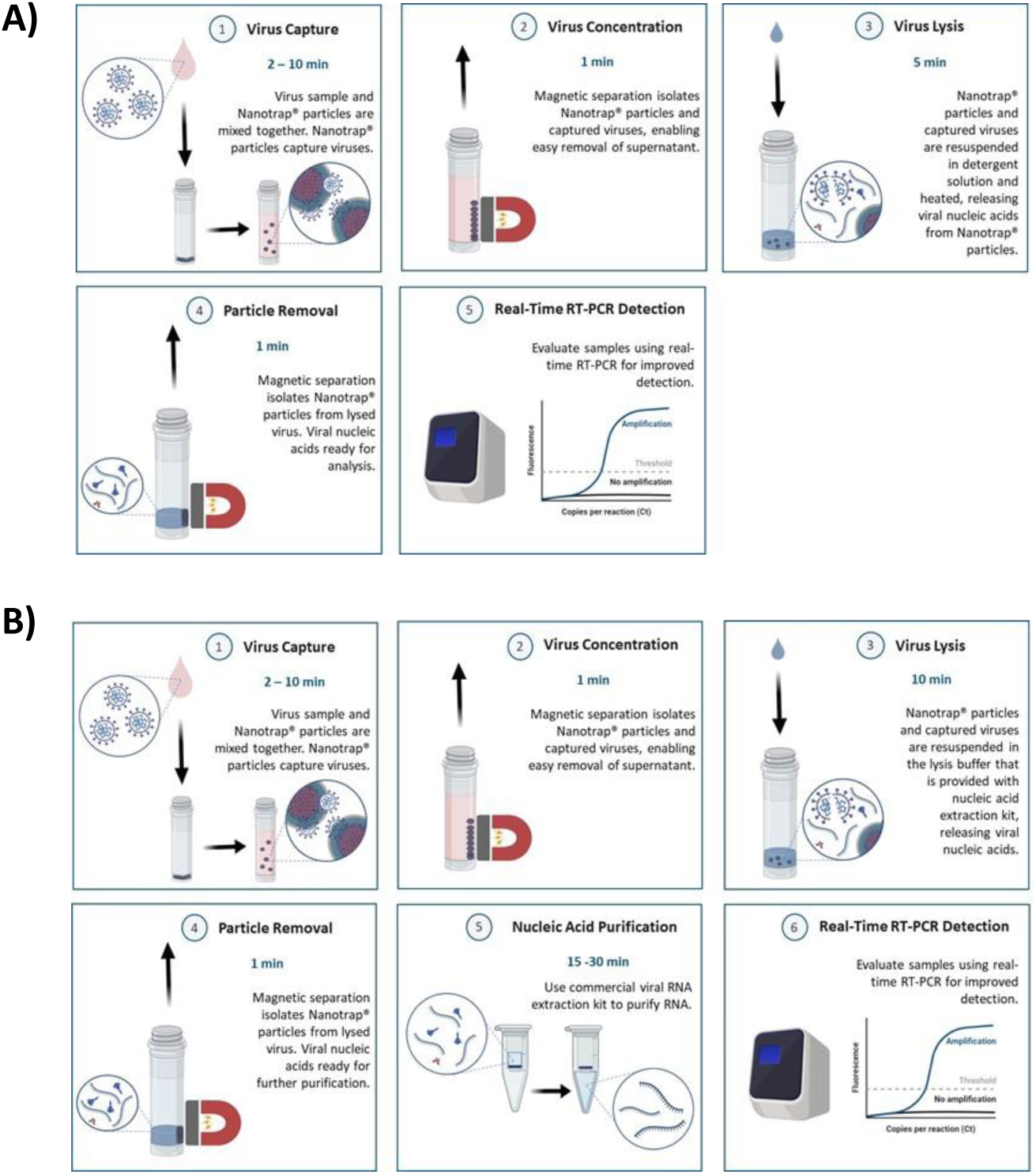
Nanotrap particles are compatible with multiple extraction methods. Sample is mixed with Nanotrap particles and incubated for 2-10 minutes. Nanotrap particles are pelleted with a magnet, the supernatant is removed, and the particle pellet is resuspended in a viral lysis solution. The Nanotrap particles are pelleted with a magnet, and (**A**) the supernatant can be used directly in real-time RT-PCR (total workflow time is 10-20 minutes) or (**B**) the supernatant can be processed with a commercial RNA extraction kit prior to viral detection (total workflow time is 30-50 minutes.)

We started by evaluating the impact that incubation time has on the amount of virus captured from a transport medium sample. We spiked 1,000,000 copies/mL of heat-inactivated SARS-CoV-2 into blank transport medium. Then we de-pooled the sample and mixed 1 mL aliquots with Nanotrap particles. The particles were allowed to incubate for either 2, 5, 10, or 30 minutes. The 2-minute and 5-minute incubation samples were not agitated during incubation. The 10-minute and 30-minute samples were inverted once every 5 minutes to keep the sample mixed, with no additional agitation. A sample without Nanotrap particles was used as a control (the 0-minute incubation time point in **Fig. 2A**). After virus capture, the particles and captured virus were concentrated using a magnet and the supernatant was removed. The pellet was resuspended in PBS and the lysis buffer from the QIAamp Viral RNA Mini Kit and incubated for 10 minutes. Viral nucleic acids were purified and analyzed using the Primerdesign assay as described in the methods section. Results in **Fig. 2A** show that virus capture is fast, as there was no significant difference in Ct values across the incubation times. Furthermore, capturing and concentrating SARS-CoV-2 from the samples prior to RNA extraction improved Ct values by about 3.

**Figure 2:**
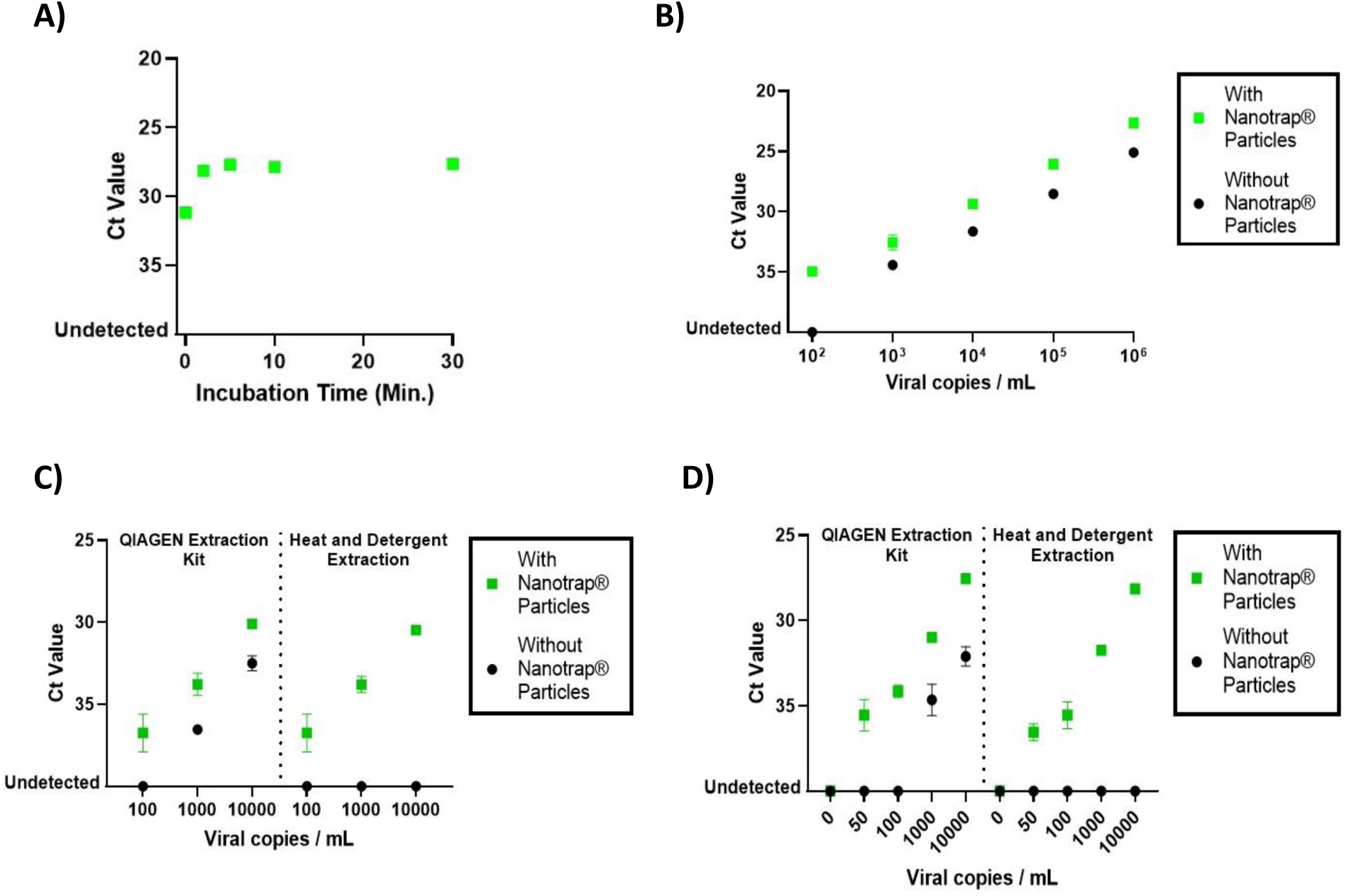
Nanotrap particles rapidly capture heat-inactivated SARS-CoV-2 in multiple sample matrices and improve detection by real-time RT-PCR. **A)** Heat-inactivated SARS-CoV-2 was spiked into transport medium at 1,000,000 copies/mL. One milliliter samples were added to Nanotrap particles and incubated for 2, 5, 10, or 30 minutes. Virus was extracted using Qiagen’s QIAamp Viral RNA Mini Kit, and real-time RT-PCR was performed using Primerdesign Ltd’s COVID-19 Genesig Real-Time CE-IVD/EUA PCR assay. A sample without Nanotrap particle processing was used to represent a 0-minute incubation with Nanotrap particles. **B)** Heat inactivated SARS-CoV-2 was spiked into viral transport medium at 100; 1,000; 10,000; 100,000; or 1,000,000 copies/mL. One milliliter samples at each viral titer were added to Nanotrap particles and incubated for 5 minutes. Virus was extracted using the QIAamp Viral RNA Mini Kit and real-time RT-PCR was performed using IDT’s 2019 nCoV CDC EUA assay. Samples at each titer without Nanotrap particles were used as controls. **C)** Heat-inactivated SARS-CoV-2 was spiked into transport medium at 100; 1,000; or 10,000 copies/mL. One milliliter samples at each viral titer were added to Nanotrap particles and incubated for 5 minutes. Virus was extracted using either the QIAamp Viral RNA Mini Kit or heat and detergent extraction. Viral detection was performed by real-time RT-PCR using the IDT 2019 nCoV CDC EUA assay. Samples without Nanotrap particles were processed by each extraction method as controls. **D)** Heat-inactivated SARS-CoV-2 was spiked into saliva at 50; 100; 1,000; or 10,000 copies/mL. Saliva aggregates were allowed to settle to the bottom of the tube for 3 minutes. The liquid saliva at each viral titer was then withdrawn, diluted as 1 volume saliva plus 2 volumes in PBS/Tween, and 1.8 mL of each dilution was added to Nanotrap particles. After an incubation of 10 minutes, the virus was extracted from the Nanotrap particles using either QIAamp Viral RNA Mini Kit or heat and detergent extraction. Viral detection was performed by real-time RT-PCR using the IDT 2019 nCoV CDC EUA assay. Samples without Nanotrap particles were processed by each extraction method as controls.

We then asked whether this method could work across a range of viral titers. Heat-inactivated SARS-CoV-2 was spiked into transport medium at 100 copies/mL; 1,000 copies/mL; 10,000 copies/mL; 100,000 copies/mL; and 1,000,000 copies/mL. One mL aliquots of each viral sample were processed with Nanotrap particles using a 5-minute capture time, followed by RNA extraction using the QIAamp Viral RNA Mini Kit. The same extraction kit also was used to extract RNA from the samples without Nanotrap particle processing. Viral RNA was then detected using real-time RT-PCR. Results in **Fig. 2B** show that capturing and concentrating SARS-CoV-2 prior to viral extraction improves Ct values across all tested viral titers. In samples containing 1,000 copies/mL of SARS-CoV-2 or more, the Ct improvement in the Nanotrap particle-processed samples was between 2 and 3. In the sample containing 100 copies/mL, the Nanotrap particles enabled detection of the virus, whereas in the absence of Nanotrap particle concentration of the virus, this sample tested negative in the real-time RT-PCR assay.

In recent months, as commercial nucleic acid extraction kits have become supply-chain limited, there have been pre-publication articles showing that it is possible to detect SARS-CoV-2 RNA from samples without an RNA extraction step ^14^. As such, we wanted to explore whether our method could be used to improve the detection of SARS-CoV-2 when used with a direct extraction method that utilizes heat and detergent. To do so, we spiked transport medium with 100 copies/mL, 1,000 copies/mL, or 10,000 copies/mL of heat-inactivated SARS-CoV-2. Samples were processed with and without Nanotrap particles, using either the QIAamp Viral RNA Mini Kit or the direct extraction method.

Results in **Fig. 2C** demonstrate that Nanotrap particles are compatible with both viral extraction methods across the range of viral titers tested and that Nanotrap particles improve Ct values across all viral titers. Interestingly, when used with Nanotrap particles, both extraction methods resulted in similar Ct values for the samples spiked at 100 copies/mL, 1,000 copies/mL, and 10,000 copies/mL. We noted that the direct extraction method without Nanotrap particles resulted in no detectable virus across the range of viral titers, which could be due to PCR inhibitors in the transport medium being loaded into the real-time RT-PCR along with the viral nucleic acid ^15^.

As swab-based sample collection is technique dependent and because there have been supply chain issues with NP swabs and transport medium, a preprint article indicated that there have been promising efforts to demonstrate that saliva is a useful sample type for SARS-CoV-2 diagnostics ^16^. Several groups reportedly have demonstrated that virus is present in saliva samples, and at least one group has already received an EUA from the FDA for a saliva-based COVID-19 test ^17^. Thus, we wanted to examine whether our method could be used to capture and concentrate SARS-CoV-2 from saliva samples.

We spiked a pooled saliva sample with varying concentrations of heat-inactivated SARS-CoV-2. We allowed aggregates in the saliva to settle to the bottom of the tube (this process took 2-3 minutes in a 5 mL saliva sample), prior to taking the supernatant and diluting it in a PBS solution containing a small amount of detergent. We then added Nanotrap particles to one milliliter of the diluted saliva sample and incubated for 10 minutes at room temperature with an inversion to mix the sample at 5 minutes. Viral extraction was performed using the QIAamp Viral RNA Mini Kit or the direct extraction method and real-time RT-PCR results were compared to samples that had not undergone Nanotrap particle processing.

The results in **Fig. 2D** demonstrate that capturing and concentrating SARS-CoV-2 from the diluted saliva sample improved real-time RT-PCR results by 3-4 Cts at viral titers of 1,000 and 10,000 copies/mL and enabled detection at 50 copies/mL and 100 copies/mL when they were otherwise undetectable. Furthermore, much like the results with transport media samples, Nanotrap particles enabled the use of the direct extraction method.

### Nanotrap^®^ Particles Capture and Concentrate Infectious SARS-CoV-2

As heat-inactivated SARS-CoV-2 is not necessarily representative of real-life testing paradigms, we assessed whether our method was compatible with infectious SARS-CoV-2. The method described above was followed, except that during the virus capture stage Nanotrap particles were incubated for 30 minutes with constant agitation. The results in **Fig. 3A** demonstrate that our method can capture and concentrate infectious SARS-CoV-2 from transport medium across multiple virus concentrations and can improve detection of that virus using a real-time RT-PCR assay.

**Figure 3:**
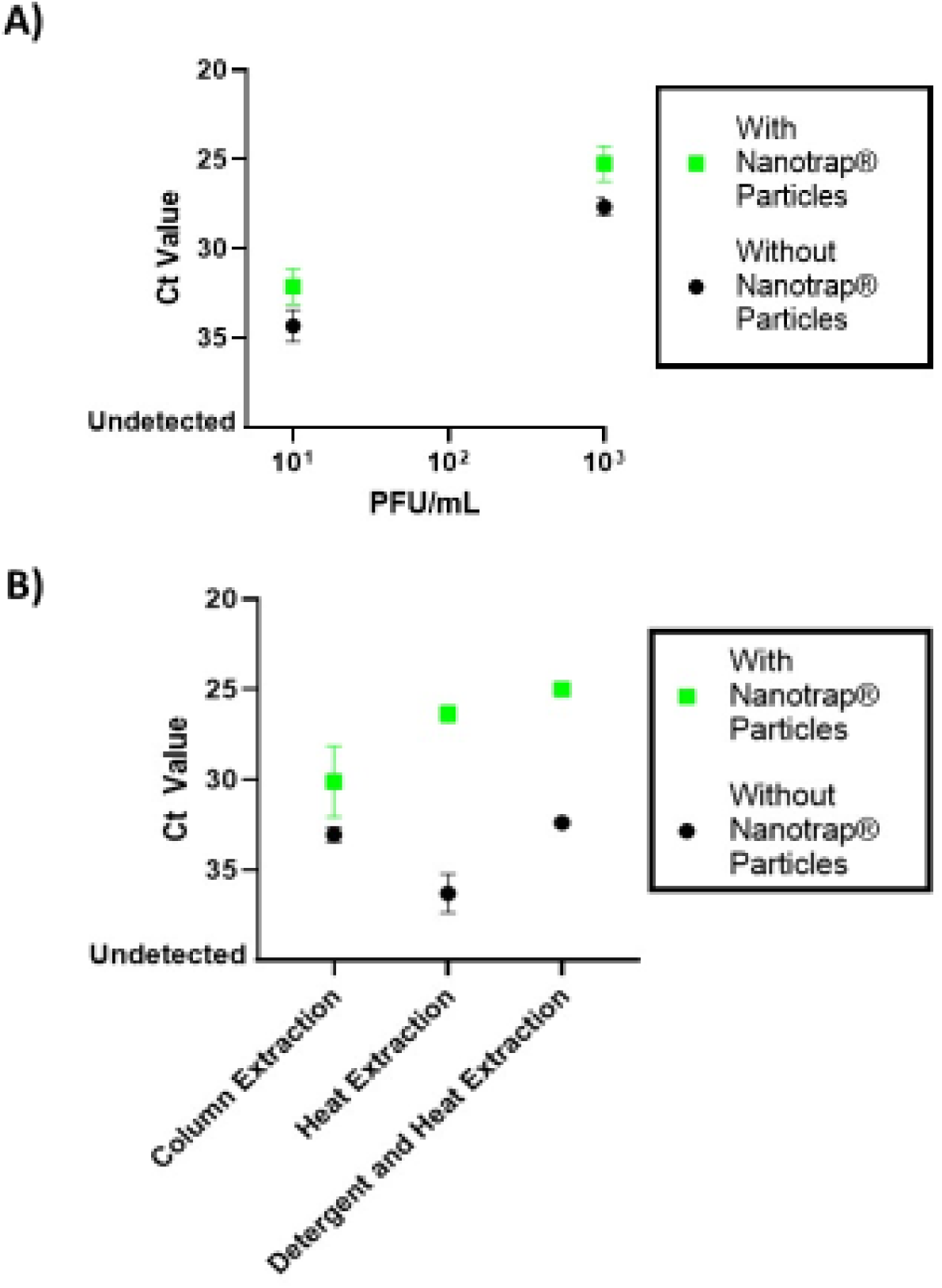
Nanotrap particles capture infectious SARS-CoV-2 from transport medium and improve detection by real-time RT-PCR. **A)** SARS-CoV-2 was spiked into viral transport medium at 10 or 1,000 pfu/mL. One milliliter samples were added to Nanotrap particles and incubated for 30 minutes. Viral extraction was performed using TRIzol LS and the RNeasy Kit, and viral detection was performed by real-time RT-PCR. Samples without Nanotrap particles were processed as controls. **B)** SARS-CoV-2 was spiked into viral transport medium at 1,000 pfu/mL. One milliliter samples were added to Nanotrap particles and incubated for 30 minutes. Viral extraction was performed using the RNeasy Kit, heat extraction, or detergent and heat extraction methods. Viral RNA was detected by real-time RT-PCR. Samples without Nanotrap particles were used as controls.

We next evaluated whether we could use Nanotrap particles with infectious virus and a direct extraction method. To accomplish this, we spiked infectious SARS-CoV-2 in 1 mL of transport medium, added Nanotrap particles, and after 30 minutes of constant agitation Nanotrap particles were pelleted with a magnet. Pelleted Nanotrap particles were either directly treated with lysis buffer followed by RNA purification on a column; or they were mixed with water and heated for 10 minutes at 95°C to release the RNA directly into the supernatant; or they were mixed with detergent and heated for 10 minutes at 95°C to release the RNA directly into the supernatant. The results in **Fig. 3B** show that the use of Nanotrap particles improves viral RNA recovery for all tested methods. Moreover, these findings are in agreement with data in **Fig. 2C** and confirm that Nanotrap particles can be used for direct SARS CoV-2 RNA purification from transport medium without the use of RNA purification kits. Interestingly, the yield of the recovered RNA for a heat and detergent method combined with Nanotrap particles is significantly higher than the Nanotrap particle method combined with the column-based RNA purification kits.

### Nanotrap^®^ Particles Can Eliminate False Negatives by Concentrating SARS-CoV-2 Prior to Testing

We next asked whether our method would improve detection of SARS-CoV-2 in samples that had been collected from patients. We obtained two sets of transport medium diagnostic remnant samples that previously had been tested for SARS-CoV-2. The first set of samples was received from Discovery Life Sciences and included 9 samples that previously had tested positive and 5 samples that previously had tested negative for SARS-CoV-2 using the Abbott RealTime SARS-CoV-2 EUA test. A second set of remnant samples was obtained from Hancock Regional Hospital in Greenwood, IN. Eight samples previously had tested positive while 27 samples previously had tested negative for SARS-CoV-2 using the Cepheid Xpert Xpress SARS-CoV-2 EUA assay. Each of these samples was processed with and without Nanotrap particles, followed by nucleic acid extraction using the QIAamp Viral RNA Mini Kit and detection by real-time RT-PCR.

Results in **Fig. 4A** and **Fig. 4B** demonstrate that our method is able to detect SARS-CoV-2 in all 17 samples that had previously tested positive. These results also demonstrate that concentrating virus from transport medium samples prior to RNA extraction significantly improves detection of SARS-CoV-2 in low viral load diagnostic remnant samples as compared to those same samples processed without Nanotrap particles, (average improvement in Ct value when using Nanotrap particles for low viral load samples is 3.1; n = 10). These improvements in Ct value are consistent with the results we saw with our contrived samples.

**Figure 4:**
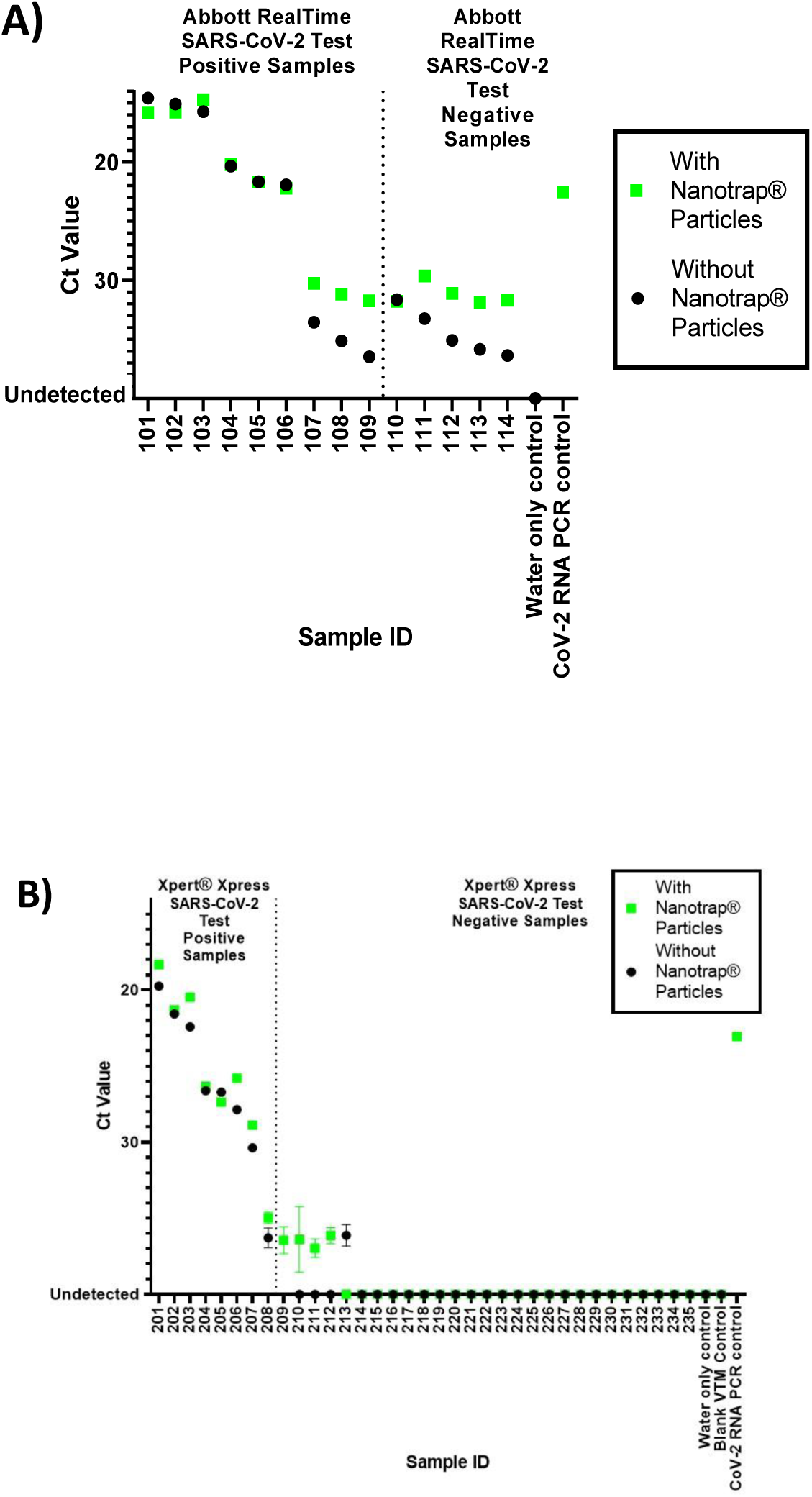
Nanotrap particles improve SARS-CoV-2 detection in low viral load diagnostic remnant samples. Forty-nine swab in transport medium diagnostic remnant samples, (**A**) 14 previously tested for SARS-CoV-2 by the Abbott RealTime SARS-CoV-2 EUA assay (101-114) and (**B**) 35 previously tested for SARS-CoV-2 by the Cepheid Xpert Xpress SARS-CoV-2 EUA assay (201-235), underwent Nanotrap particle processing. One milliliter of each sample was added to Nanotrap particles and incubated for 5 minutes. Viral extraction was performed using the QIAamp Viral RNA Mini Kit and viral RNA was detected using real-time RT-PCR. Samples 101-109 and Samples 201-208 had previously tested positive for SARS-CoV-2 on the Abbott and Cepheid assays, respectively. Samples 110-114 and Samples 210-235 had previously tested negative. Each diagnostic remnant sample also underwent processing without Nanotrap particles for equivalency with previous results. An uninfected transport medium sample was used as a negative control.

Of even greater interest to us were the results that suggest that using Nanotrap particles to concentrate the virus from the transport medium prior to nucleic acid extraction can identify potential false negatives. In four of the samples that previously had been reported as negative for SARS-CoV-2 (Samples 209-2012 in **Fig. 4B**), we observed Ct values that were indicative of low levels of virus when we used our method. Note that for Sample 213, Nanotrap particle processing did not result in detection of SARS-CoV-2 even though SARS-CoV-2 was detected in the sample without Nanotrap particle processing (i.e. with QIAamp Viral RNA Mini Kit extraction only). This could be due to sample storage conditions; these samples had been subjected to at least one freeze-thaw cycle, which may have lysed virus in the transport medium prior to the addition of the Nanotrap particles. The Nanotrap particles can capture whole virions but cannot capture free-floating nucleic acids, whereas we assume that a QIAamp Viral RNA Mini Kit extraction kit would capture and purify free-floating viral nucleic acids, regardless of whether those came from previously lysed or recently lysed viruses. No SARS-CoV-2 was observed in our negative control (water), indicating the presence of SARS-CoV-2 in the five previously negative samples is not due to PCR background.

We also noted that the Nanotrap particles did not consistently improve the Ct values of the high viral titer samples (see, for example, Samples 101-106 in **Fig. 4A**). These results suggest that the Nanotrap particles may become saturated at very high viral titers. This could be addressed by using a larger amount of Nanotrap particles in sample processing, though that may have implications for the very low viral titer samples. For immediate testing ramp-up needs, however, this method appears to be very useful, as getting a very accurate assessment of the viral load is much less clinically relevant than knowing whether virus is present in a sample.

### Nanotrap^®^ Particles Capture and Concentrate SARS-CoV-2 from Large Sample Volumes

The diagnostic industry is racing to increase testing capabilities for COVID-19. Despite the impressive efforts thus far, we are still far short of the total number of daily tests that public health experts suggest are necessary ^18-20^. To address this shortage, the U.S. Department of Health and Human Services has called for the development of new diagnostic tools, including “pooling of samples from multiple (5, 10, or 20) individuals in a single test” in circumstances where the overall prevalence of the disease is low ^21^.

Others have demonstrated that pooling of samples for testing for SARS-CoV-2 can increase test capacity but that it would introduce a risk of borderline positive samples escaping detection ^22^. Therefore, we asked whether Nanotrap particles could capture infectious SARS-CoV-2 from a large volume of a highly diluted sample, mimicking sample pooling prior to viral extraction, without sacrificing detection of low viral load samples. Infectious SARS-CoV-2 was spiked into 1 mL of transport medium at 100 pfu/mL. One hundred microliters of diluted virus were used for RNA extraction and purification with RNeasy kit. The rest of the diluted virus was mixed with 4 or 9 mL of uninfected transport medium. Nanotrap particles were incubated with the samples for thirty minutes with constant agitation to capture the virus and viral extraction was achieved using direct extraction method. Viral detection was performed using real-time RT-PCR. Results in **Fig. 5A** indicate that Nanotrap particle processing dramatically improves Ct values in the diluted samples (by 5-6 Ct values) compared to the sample that was not processed with Nanotrap particles. Furthermore, the difference of about 1 Ct value between the 5 mL and 10 mL samples that underwent Nanotrap particle processing is consistent with the notion that two times as much virus had been spiked into the 10 mL sample.

**Figure 5:**
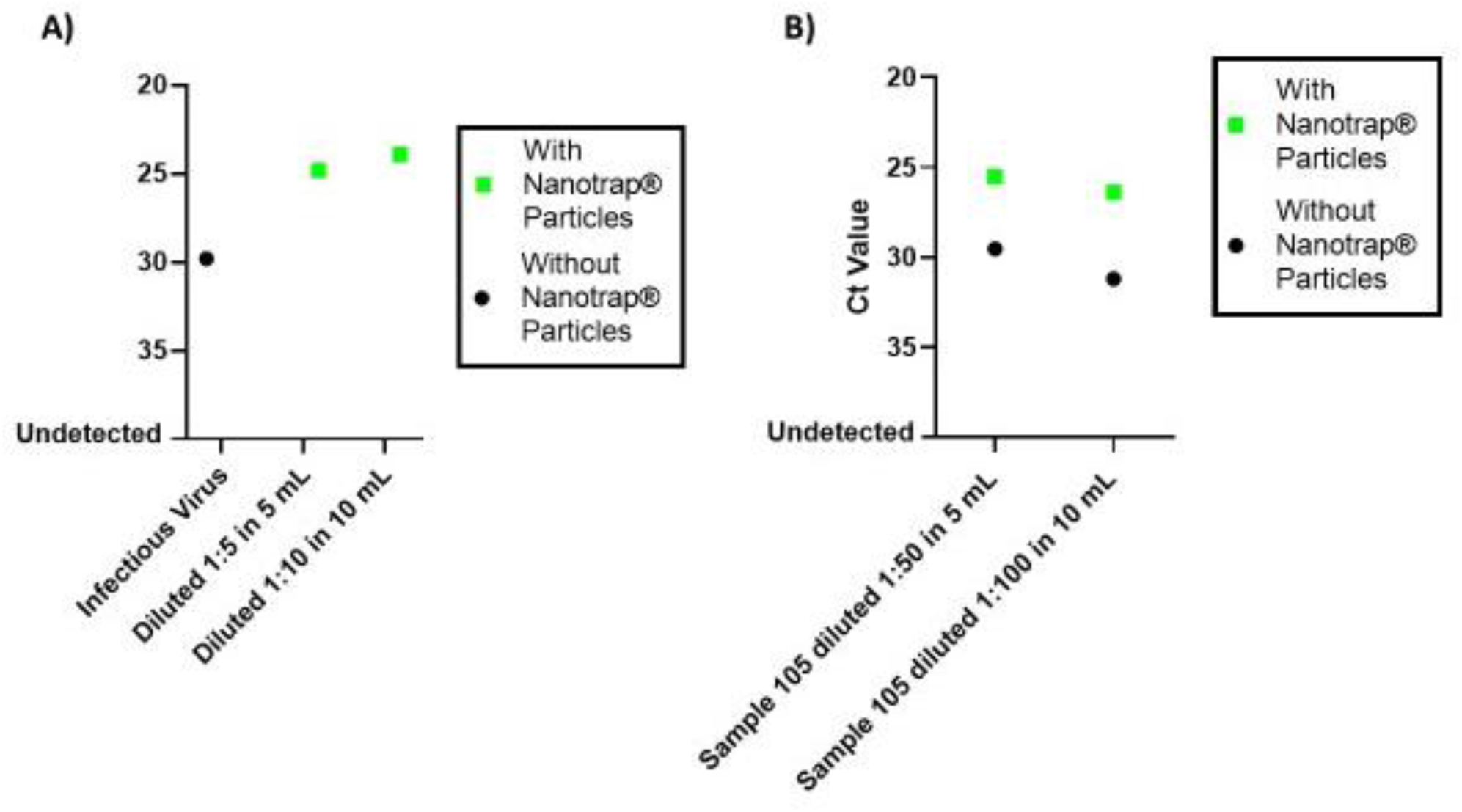
Nanotrap particles improve detection of SARS-CoV-2 from pooled sample mimics. **A)** Infectious SARS-CoV-2 was spiked into 1 mL of transport medium at 100 pfu/mL. Virus spiked-medium was mixed with 4 or 9 mL of uninfected transport medium in the presence or absence of Nanotrap particles. The samples without Nanotrap particles were processed using the RNeasy kit. Samples with Nanotrap particles were incubated for 30 minutes prior to heat and detergent extraction. Viral RNA was detected using real-time RT-PCR. **B)** One hundred microliters of SARS-CoV-2-infected viral transport medium from Sample 105 was spiked into uninfected viral transport medium to a total volume of 5 mL or 10 mL. Nanotrap particles were added to the samples and incubated for 5 minutes. Viral extraction was performed using the QIAamp Viral RNA Mini Kit and SARS-CoV-2 detection was performed by real-time RT-PCR. Samples without Nanotrap particles were processed as controls.

To determine whether our method would work for a larger sample volume with a more realistic diagnostic sample, we diluted 100 µL of diagnostic remnant Sample 105 into uninfected transport medium at 1:50 and 1:100 dilutions (5 mL and 10 mL total volume); each diluted sample contained the same total amount of virus. We then added Nanotrap particles to the sample and allowed them to incubate for five minutes with no agitation. Viral extraction was performed using the QIAamp Viral RNA Mini Kit. The 5 mL and 10 mL samples were also processed without Nanotrap particles to serve as controls. The results in **Fig. 5B** show that using Nanotrap particles to capture and concentrate SARS-CoV-2 from 5 mL and 10 mL transport medium samples significantly improves detection of the virus. Interestingly, the Nanotrap particles were able to recover roughly the same amount of virus in both the 5 mL and 10 mL dilutions, suggesting that the full amount of virus was recovered in both samples, and that even larger sample volumes could be processed using the same method. Collectively, these data indicate that our method could enable much better detection of SARS-CoV-2 in samples that have been pooled prior to RNA extraction. This suggests a path toward testing large numbers of patient pooled samples for presence of SARS-CoV-2 while reducing the reagents and labor needed to perform the nucleic acid extraction and detection steps.

## Discussion

The simple method we have developed can take as little as 10 minutes to prepare SARS-CoV-2 RNA for downstream testing and obviates the need for commercial nucleic acid extraction kits. Using contrived samples with heat-inactivated and infectious SARS-CoV-2, we have demonstrated significant improvements in Ct values from transport medium and saliva samples with both commercial nucleic acid extraction kits and a direct extraction method.

Contrived samples lack the biological complexity (i.e. cells, cell debris, bacteria) contained in patient samples. Using 49 diagnostic remnant samples, we demonstrated that our method can improve detection of SARS-CoV-2 in real samples and eliminate false negatives. Our method had 100% concordance with all of the samples that previously tested positive, improved the real-time RT-PCR signal by an average of 3.1 Ct values in the low viral load samples, and identified SARS-CoV-2 in four of the samples that had previously tested negative.

The national and international targets for numbers of daily tests are astronomical. There is a growing consensus that pooling of samples is going to be one of the methods that must be implemented in order to reach those numbers of daily tests, an opinion which is shared by the government of the United States ^22, 23^. Our method was able to concentrate infectious SARS-CoV-2 from contrived samples and SARS-CoV-2 from diagnostic remnant samples in pooled patient sample mimics, an approach which is a promising way forward to address the massive testing scale-ups that are necessary.

There are several limitations to this study. First, we only had fourteen diagnostic remnant samples that had been previously tested on the Abbott RealTime SARS-CoV-2 EUA assay, and only five of those samples had previously tested negative for SARS-CoV-2. A larger set of samples would help us assess how our method compares to that assay. Second, for all of the diagnostic remnant samples that we used, the storage conditions were inconsistent. Some samples were unintentionally subjected to multiple freeze-thaw cycles, others were only subjected to one free-thaw cycle, and others were shipped and stored at 4°C and never frozen. Third, it is important to note that we used a different detection assay (from IDT) than had been used previously on the diagnostic remnant samples; therefore, we cannot make any direct conclusions regarding whether Nanotrap particles would improve the performance of those assays. Additionally, our saliva data is limited to contrived samples and it is therefore difficult to gauge how effective our method would be for saliva SARS-CoV-2 testing in a clinical setting.

Despite these limitations, we are confident that the method described here can offer increased sensitivity for SARS-CoV-2 molecular testing from transport medium and saliva while enabling the use of faster and easier nucleic acid extraction methods, which will help address supply chain limitations. Also, as we have previously demonstrated that the Nanotrap particles used in this study can also capture and concentrate influenza viruses, RSV, and other coronaviruses ^13^, we are confident that the method described here will be compatible with the respiratory virus panel tests that are under development. Moreover, while we did not address assays for viral antigens, next generation sequencing assays, or antibody assays in this work, Nanotrap particles can be used to improve those testing modalities ^10, 11, 24-26^ and we look forward to expanding our work into that area.

## Acknowledgements

The following reagent was deposited by the Centers for Disease Control and prevention and obtained through BEI Resources, NIAID, NIH, SARS-Related Coronavirus 2, Isolate USA WA1/2020, NR-52281. The following reagent was deposited by the Centers for Disease Control and Prevention and obtained through BEI Resources, NIAID, NIH: SARS-Related Coronavirus 2, Isolate USA-WA1/2020, Heat Inactivated, NR-52286. Funding provided, in part, by Schmidt Futures.

## Author Contributions

R.A.B., I.A., A.P., J.W., D.M., P.A., C.L., V.C., S.B., B.K., R.K., and D.G. carried out the experiments, contributing to cell culture, viral capture, viral extraction, and viral detection. R.R., S.M., S.S., O.S., A.M., and S.P. contributed to the production and quality control of the Nanotrap^®^ particles. R.B., E.P., T.J., and R.D. contributed to data presentation and formatting. R.A.B., I.A., A.P. analyzed and interpreted the data. R.A.B., I.A., A.P., R.B., K.K.H., and B.L. were involved in experimental design and the writing and editing of the manuscript while also providing the overall direction and coordination of this study. All authors approved the manuscript.

